# Antagonism Rather Than Synergy: Ibuprofen-Antifungal Interactions Depend on Strain Genetics and Nutrient Environment

**DOI:** 10.64898/2026.01.22.701080

**Authors:** Amrish Prabakaran, Himanshu Sinha

## Abstract

Drug interaction outcomes-synergism, additivity, or antagonism-represent complex phenotypes. While drug repurposing aims to identify compounds that potentiate conventional antifungals, when given in combination, the modulators of these interactions in fungi remain largely unexplored. We hypothesize that the response to repurposed-conventional antifungal pairs is a complex trait modulated by genetic and environmental factors.

To study the impact of genotype on the outcome, we screened six diverse *Saccharomyces cerevisiae* isolates, including clinical, wild, and fermentation strains, for their responses to combinations of ibuprofen with either clotrimazole or caspofungin. We evaluated the role of the environment using rich and minimal media and assessed the influence of assay type by comparing solid- and liquid-rich media assays. Our results reveal that ibuprofen-clotrimazole interactions are highly dynamic, predominantly antagonistic, with limited synergy observed. These outcomes are significantly modulated by genetic background, media composition, assay type, and, in specific genotypes, even by the drug dosage, reflecting a complex, multi-parametric phenotype.

However, the ibuprofen-caspofungin combination is more predictable, exhibiting only synergy or additivity. Interaction outcomes correlate with baseline sensitivity to caspofungin: caspofungin-resistant isolates consistently demonstrate synergy, while sensitive strains exhibit additivity.

These findings shift the paradigm of drug discovery by demonstrating that synergism and antagonism are not static properties of drug pairs but are dynamic, context-dependent outcomes. This study highlights the need to use clinically relevant models and patient-specific isolates before clinical application, as drug interactions cannot be generalized from a single dosage, strain, or environmental condition.

## 1. Introduction

While single-drug responses are inherently complex, strains’ responses to drug combinations exhibit even greater variability, making it a complex phenotypic trait. The response to a drug combination can be broadly classified as synergy, antagonism, or additivity (Duarte & Vale, 2022). These drug-drug interaction patterns are often viewed as fixed properties of the drug compounds themselves. However, emerging studies in cancer and tuberculosis suggest that these interactions are highly context-dependent phenotypes (Chung & Chandrasekaran, 2022; Flobak et al., 2019; Goossens et al., 2020; Iorio et al., 2016; Jaaks et al., 2022; Ramón-García et al., 2011). Bacterial studies have established that drug-drug interactions are modulated by the underlying genetic architecture and environmental conditions (Brochado et al., 2018; Davis et al., 2024). With only three major antifungal classes, the focus of current antifungal drug research has shifted towards repurposing existing drugs to potentiate conventional therapies (Alabi et al., 2023; Stenkiewicz-Witeska & Ene, 2023; Xiong et al., 2025). Given that *de novo* antifungal development is slow and expensive, the strategy of using repurposed compounds as potentiators or to target resistance mechanisms could be ideal for combination therapy.

While previous research has revealed strain-specific responses to repurposed antifungal combinations in pathogenic fungi such as *Candida albicans* and *Aspergillus fumigatus*, these variations are often documented without further investigation into their underlying drivers (Król et al., 2018), hindering the clinical translation of *in vitro* synergies. Recent studies investigating novel antifungal compounds or synergistic drug pairs have predominantly focused on a limited selection of clinical isolates (Burns et al., 2024; Lefranc et al., 2024; Liang et al., 2025). While these studies occasionally include representative strains from different pathogenic species, they often overlook the significant intra-species variation that can dictate drug interaction outcomes (Pinder et al., 2025; Tancer et al., 2024). This gap arises from treating drug interactions as static properties rather than as dynamic outcomes modulated by genetics and microenvironments. The present study addresses this by screening a genetically diverse *Saccharomyces cerevisiae* collection to elucidate how intra-species variation and nutrient availability influence drug-drug interaction landscapes.

*S. cerevisiae*, an established model for dissecting complex phenotypes, shares conserved pathways with pathogenic fungi, including ergosterol biosynthesis and cell wall integrity, targeted by major antifungals (Demuyser & Van Dijck, 2019). To address the gap and dissect the influence of genetic background on drug-drug interactions, a genetically diverse subset of strains from the Saccharomyces Genome Resequencing Project (SGRP) (Liti et al., 2009) was selected. With a reported nucleotide diversity of 0.0056, this collection reflects significant intra-species variation, as the constituent strains were isolated from a wide range of ecological niches, including clinical, soil, nectar, baking, and fermentation environments. The six strains chosen for this study comprise two clinical, two fermentation, and two wild isolates adapted to distinct ecological niches. These localized ecological adaptations have likely led to extensive genetic divergence and metabolic network reorganization, potentially modulating the observed variations in drug response. Further, to evaluate the impact of the environment on modulating drug-drug interaction patterns, we used two distinct experimental formats: solid agar and liquid broth assays. These formats differ significantly in oxygen availability, drug exposure kinetics, and metabolic heterogeneity, all of which can alter strain-specific responses (Hachicho et al., 2017; Imanaka et al., 2010). Furthermore, within the solid assay framework, interactions were tested in both nutrient-rich and minimal media to elucidate the impact of nutrient availability, a factor previously shown by Davis et al. (2024) to significantly shift drug interaction outcomes.

Clotrimazole (azole class) and caspofungin (echinocandin class) were selected as representative conventional antifungals. For the repurposed drug component, we utilized the non-steroidal anti-inflammatory drug (NSAID) class, which has attracted considerable attention as a promising candidate for antifungal therapy (Babaei et al., 2024). NSAIDs exhibit antifungal activity across diverse fungal species and potentiate conventional therapies. These compounds are reported to impair biofilm formation and attenuate essential virulence factors in pathogenic fungi (Babaei et al., 2024). Within this class, ibuprofen was prioritized due to its established synergistic potential when combined with azoles (Babaei et al., 2024; Pina-Vaz et al., 2000; Ricardo et al., 2009; Rossato et al., 2021). Previous reports indicated that ibuprofen reverses azole resistance mediated by efflux pumps in *Candida spp*. (Pina-Vaz et al., 2005; Ricardo et al., 2009). The current study contrasted ibuprofen-azole interactions with those involving a mechanistically distinct antifungal class (echinocandins) to evaluate the generalizability of these interaction outcomes. Since caspofungin targets β-1,3-glucan synthase in the cell wall (distinct from efflux pump targets) (Robbins et al., 2017), ibuprofen-caspofungin synergy via efflux inhibition is mechanistically improbable, suggesting alternative pathways. Research into such non-azole combinations is sparse, with existing studies limited to specific pathogens such as *Trichosporon asahii* rather than common fungal pathogens (Yang et al., 2016). By exploring these diverse pairings, this approach addresses whether ibuprofen potentiates antifungal efficacy through a singular, conserved mechanism or via multiple, yet-to-be-characterized pathways.

Our primary findings reveal genotype as the key determinant of a strain’s response to drug combinations. Further experiments on the role of the environment showed that, for the ibuprofen-caspofungin combination, the response was primarily determined by genotype. However, for the ibuprofen-clotrimazole combination, the outcome was influenced by multiple factors, including genotype, media, assay type, and even drug concentration. These results underscore the varying impact of genetic and environmental factors on different drug combinations, highlighting the need for a more comprehensive understanding of this complex phenotype to facilitate the development of personalized treatment regimens tailored to the specific genotype of the infecting strain.

## 2. Results

### 2.1. Single drug response to antifungal and non-antifungal drugs

All six yeast strains were tested for their susceptibility to clotrimazole at 0.25, 0.50, and 1 mg/L in YPD medium (Fig. 1A). At all three concentrations, the clinical strain YJM789 exhibited uniform growth. This indicated that at both 0.25 mg/L (low) and 1 mg/L (high), the strain showed similar susceptibility to the drug and was not entirely resistant; however, its sensitivity did not increase linearly with increasing drug concentration. Similarly, the other clinical strain, 322134S, showed uniform growth across the three drug concentrations, with slightly higher resistance than YJM789. The strain NCYC demonstrated increasing susceptibility as the drug concentration increased from 0.25 to 1 mg/L. The strain DBVPG was the most resistant to clotrimazole, with resistance higher than that of any clinical isolate. However, similar to the clinical strains, it exhibited a non-linear response, with the least susceptibility at 0.25 mg/L, but was equally susceptible at both 0.50 and 1 mg/L. The Y9 strain also showed a linear increase in susceptibility with increasing drug concentration. The UWOP strain was most sensitive to clotrimazole, showing increasing susceptibility with increasing drug concentration, with no growth at 1 mg/L. We chose 0.5 mg/L of clotrimazole for the drug combination experiments to ensure all strains grew in the further phenotyping experiments.

**Figure 1:**
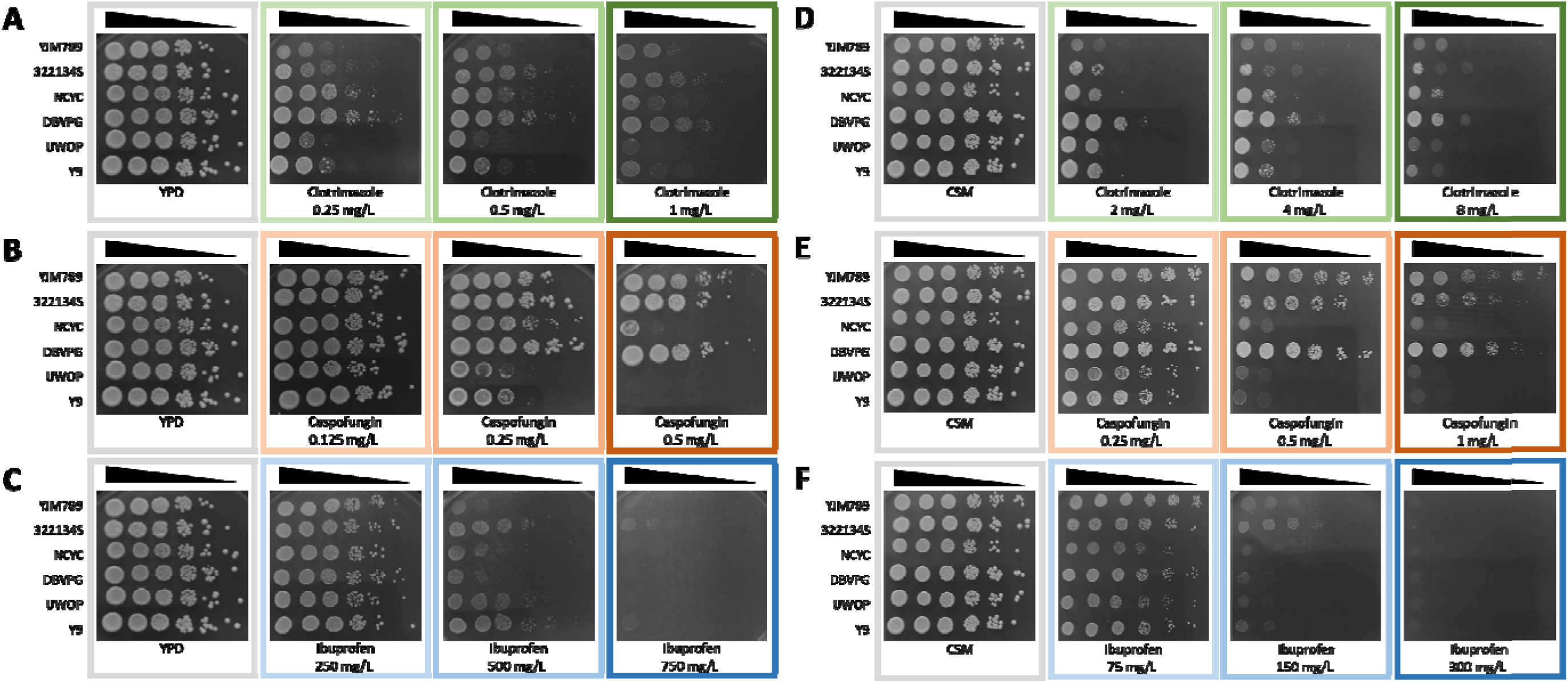
Single-drug susceptibility assays for S. cerevisiae in YPD and CSM media. Representative spot dilution images illustrate antifungal activity across six strains in YPD (A-C) and CSM (D-F) media. Drug concentrations tested include: clotrimazole at (A) 0.25, 0.5, and 1 mg/L and (D) 2, 4, and 8 mg/L; caspofungin at (B) 0.125, 0.25, and 0.5 mg/L and (E) 0.25, 0.5, and 1 mg/L; and ibuprofen at (C) 250, 500, and 750 mg/L and (F) 75, 150, and 300 mg/L.

To test the yeast strains with caspofungin in YPD, the selected concentrations were 0.125, 0.25, and 0.50 mg/L (Fig. 1B). The clinical strains YJM789 and 322134S, along with the wild-type strain DBVPG, showed high resistance to caspofungin, with minimal susceptibility even at 0.5 mg/L. In contrast, strain NCYC displayed intermediate sensitivity; it remained resistant at lower concentrations but became highly susceptible at 0.5 mg/L. Strains UWOP and Y9 showed a linear, dose-dependent increase in susceptibility and were characterized as highly sensitive to caspofungin across the tested range. Based on these profiles, a caspofungin concentration of 0.25 mg/L was selected for subsequent drug combination experiments. This concentration was used to ensure sufficient growth across all strains while allowing distinct phenotypic responses to the combination to be observed.

For ibuprofen, the strains were tested at concentrations of 250, 500, and 750 mg/L in YPD (Fig. 1C). None of the strains showed any growth defect at 250 mg/L of ibuprofen; however, it was observed that colony sizes were qualitatively smaller than those of the control medium without drug, suggesting that ibuprofen might affect metabolism, reducing the colony size. At 500 mg/L of ibuprofen, distinct phenotypic variation emerged among the isolates. Strains 322134S, UWOP, and Y9 demonstrated relative resistance, whereas YJM789, NCYC, and DBVPG were comparatively susceptible. Conversely, all strains were completely inhibited at 750 mg/L, with no visible growth recorded. Although 500 mg/L revealed phenotypic diversity, 250 mg/L was selected as the working concentration for subsequent drug combination experiments. This suboptimal concentration was chosen to ensure that ibuprofen alone did not cause significant growth defects, thereby allowing clear observation of its potential to potentiate the antifungal activity of clotrimazole or caspofungin.

In the CSM medium, the selected clotrimazole concentrations for testing were 2, 4, and 8 mg/L (Fig. 1D). At 2 mg/L, most strains showed uniform susceptibility to clotrimazole, except for DBVPG, which showed slight resistance. This general pattern of susceptibility persisted at higher concentrations. Consequently, a clotrimazole concentration of 2 mg/L was selected for the drug-combination experiments, as it provided sufficient phenotypic variation among the isolates.

Caspofungin was tested in CSM at 0.25, 0.5, and 1 mg/L (Fig. 1E). Consistent with observations in YPD media, strains YJM789, 322134S, and DBVPG exhibited high resistance to caspofungin in CSM, with increasing susceptibility at higher drug concentrations. Conversely, the strains NCYC, UWOP, and Y9 were highly sensitive to the caspofungin. At concentrations of 0.5 and 1 mg/L, these strains exhibited pronounced growth defects. Based on these profiles, a caspofungin concentration of 0.5 mg/L was selected for subsequent drug combination experiments in CSM media, as it provided an optimal baseline for evaluating potential synergistic or antagonistic interactions.

Yeast strains were tested with ibuprofen in CSM at the concentrations of 75, 150, and 300 mg/L (Fig. 1F). At 75 mg/L, all strains remained viable across the entire dilution series. Consistent with previous observations in YPD media, ibuprofen appeared to reduce colony size relative to the control medium without drug, suggesting a sub-inhibitory impact on growth rate. At 150 mg/L, severe growth defects were observed. Clinical strains YJM789 and 322134S exhibited relative resistance and maintained marginal growth, while the remaining strains failed to grow in any dilution. At the highest concentration of 300 mg/L, none of the strains grew. For subsequent drug combination experiments, a suboptimal concentration of 75 mg/L was selected, as it avoided severe growth defects while allowing detection of potential potentiation effects of the repurposed drug ibuprofen.

Broth dilution assays were performed to determine the minimum inhibitory concentrations (MICs) of various drugs across six strains, and checkerboard assays were performed to cover the full concentration range from 0 to the respective MICs. The MIC values for each drug-strain combination are summarized in Table 2. For clotrimazole, five strains had an MIC of 8 mg/L, while strain 322134S had a higher MIC value of 16 mg/L, likely due to its flocculation phenotype. The MICs for caspofungin varied, with YJM789, 322134S, and DBVPG showing higher values compared to the other strains, in line with observations from solid agar assays. The caspofungin-sensitive strains NCYC, UWOP, and Y9 exhibited lower MIC values of 0.5, 0.125, and 0.25 mg/L, respectively. For the non-antifungal drug ibuprofen, the MIC was consistently 1,000 mg/L across all strains in the study.

**Table 1:** Six *S. cerevisiae* strains and their sources.

| Strains | Geographic | Source |
| --- | --- | --- |
| YJM789 | USA, North America | Clinical (Lung) |
| 322143S | UK, Europe | Clinical (Throat sputum) |
| NCYC110 | West Africa, Africa | Fermentation (Ginger beer) |
| DBVPG1373 | Netherlands, Europe | Wild (Soil) |
| UWOPS03-461.4 | Malaysia, Asia | Wild (Nectar, Bertram palm) |
| Y9 | Indonesia, Asia | Fermentation (Ragi, similar to sake wine) |

**Table 2:**
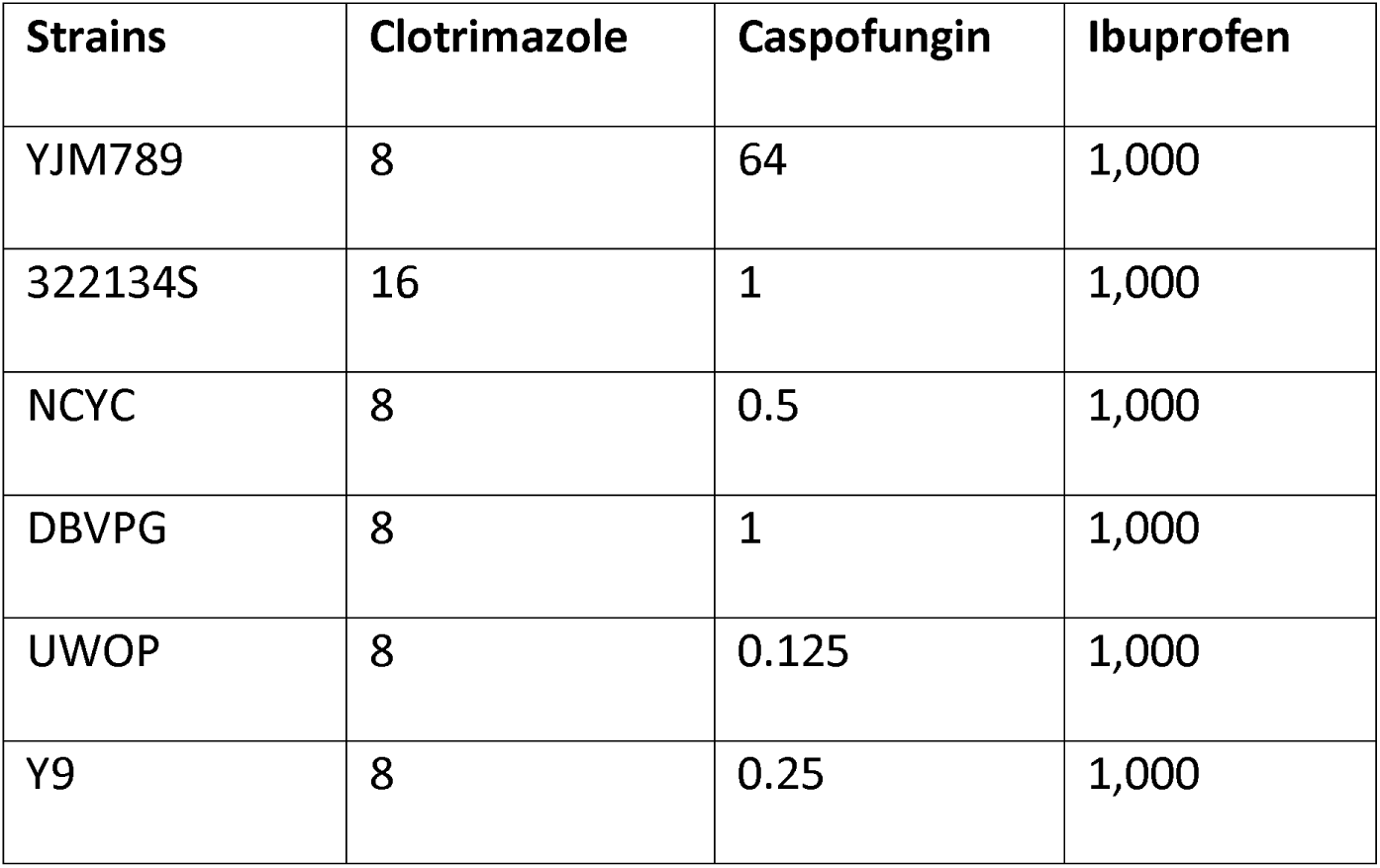
Checkerboard MICs of strains against individual drugs (mg/L).

| Strains | Clotrimazole | Caspofungin | Ibuprofen |
| --- | --- | --- | --- |
| YJM789 | 8 | 64 | 1,000 |
| 322134S | 16 | 1 | 1,000 |
| NCYC | 8 | 0.5 | 1,000 |
| DBVPG | 8 | 1 | 1,000 |
| UWOP | 8 | 0.125 | 1,000 |
| Y9 | 8 | 0.25 | 1,000 |

### 2.2. Ibuprofen-clotrimazole combination: interdependence of genotype and media composition

We investigated the effect of adding the non-antifungal drug, ibuprofen, to the antifungal drugs clotrimazole or caspofungin. Drug interaction patterns were determined by comparing the growth profiles of strains on plates containing the drug combination to those on plates containing the antifungal drug alone. A response was classified as additive (indifferent) when no observable change in growth occurred between the combination and the antifungal alone, suggesting that ibuprofen did not modulate antifungal activity. Synergy and antagonism were defined as marked reductions or increases in growth, respectively, in the presence of the drug combination relative to the antifungal control. Specifically, synergy indicated potentiation of the antifungal effect, whereas antagonism indicated attenuation of drug efficacy.

In YPD media, the strain YJM789 exhibited a 100-fold reduction in growth when exposed to the drug combination compared with antifungal monotherapy (Fig. 2A), indicating a synergistic interaction between the drugs.

**Figure 2:**
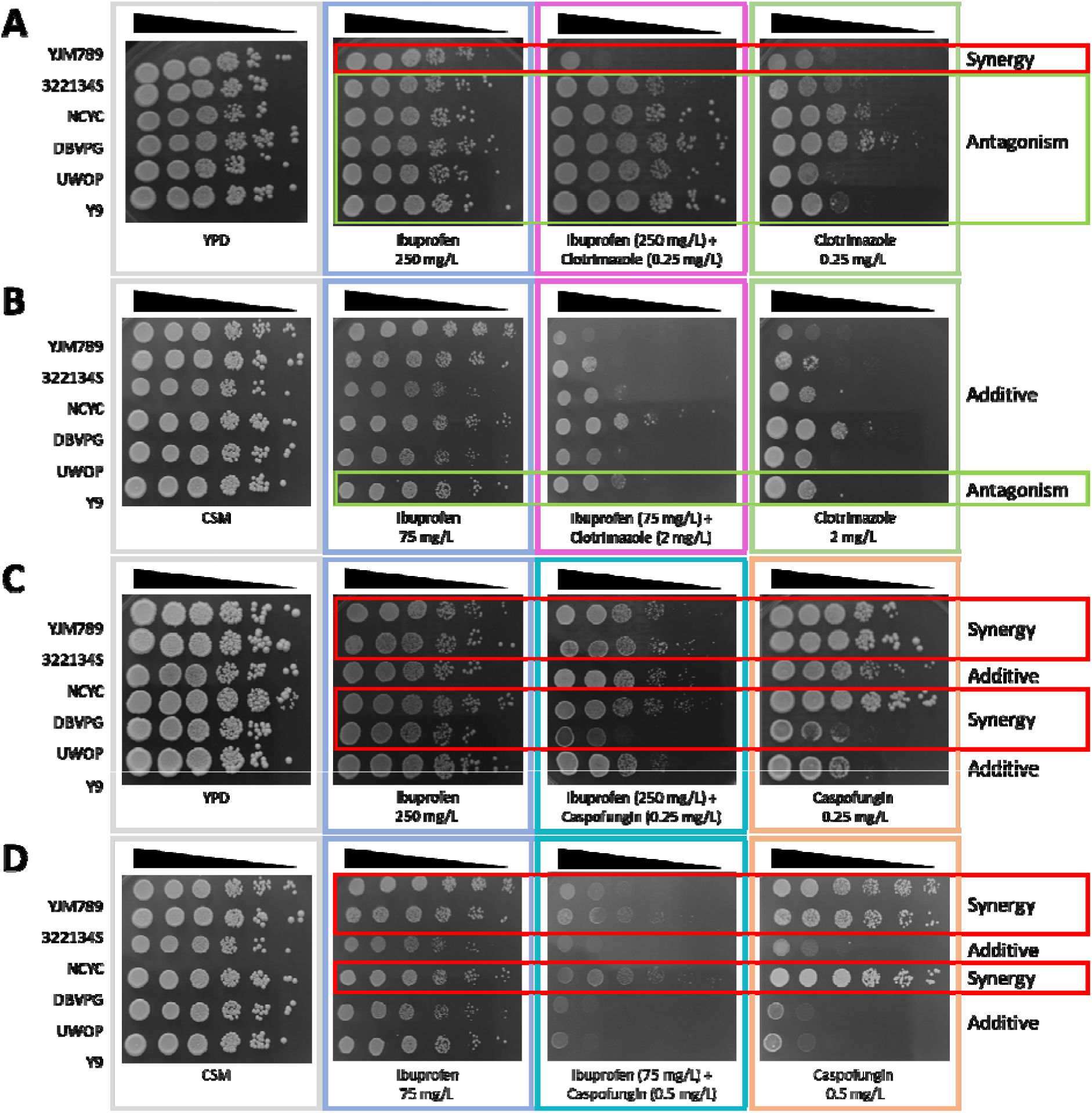
Spot dilution assays of S. cerevisiae drug combinations in YPD and CSM media. Representative images demonstrate strain performance after 48 hours of incubation at 30°C in YPD (A, B) and CSM (C, D) media. Each panel compares a combination treatment to its respective single drug treatment and a no-drug control. Specific concentrations evaluated are as follows. In YPD media: (A) Ibuprofen (250 mg/L) + Clotrimazole (0.25 mg/L); (B) Ibuprofen (250 mg/L) + Caspofungin (0.25 mg/L). In CSM media: (C) Ibuprofen (75 mg/L) + Clotrimazole (2 mg/L); (D) Ibuprofen (75 mg/L) + Caspofungin (0.5 mg/L). Red and green boxes indicate specific strains in which the combination treatment results in synergistic or antagonistic growth, respectively, compared to the single drug.

On the contrary, the other clinical strain, 322134S, demonstrated a 10-fold increase in growth in the presence of the combination, suggesting antagonistic behavior of the drugs. The remaining strains displayed 10- to 10,000-fold increased growth on the drug combination plates, indicating that the drugs acted antagonistically across those genetic backgrounds as well. In summary, in the YPD medium, ibuprofen potentiated the activity of clotrimazole in a single strain (YJM789), while it antagonized the efficacy of clotrimazole in all other tested strains.

When the ibuprofen-clotrimazole combination was evaluated in CSM media (Fig. 2B), strain Y9 exhibited a 10-fold increase in growth, suggesting an antagonistic interaction of the drugs. In contrast, the remaining strains showed no significant growth alterations compared with clotrimazole monotherapy, indicating an additive interaction between the drug combination.

Strain YJM789 exhibited a synergistic interaction between ibuprofen and clotrimazole in YPD medium, which transitioned to an additive interaction in CSM medium. On the other hand, strain Y9 demonstrated persistent antagonism for the drug pair across both media types. Strains 322134S, NCYC, DBVPG, and UWOP, which revealed antagonistic interaction in YPD, exhibited additive interactions of the drugs when tested in CSM. These significant shifts in interaction profiles for the same strains and drug combinations underscore the influence of the nutritional environment on pharmacological outcomes. Collectively, these findings suggest that the response to the ibuprofen-clotrimazole combination is a complex phenotype modulated by both the genetic background of the strain and specific nutritional (environmental) conditions.

### 2.3. Dominance of strain genotype in determining ibuprofen-caspofungin interaction profile

When the ibuprofen-caspofungin combination was evaluated in YPD media (Fig. 2C), clinical strains YJM789 and 322134S, along with strains DBVPG and UWOP, exhibited a 10-fold reduction in growth. This indicated a synergistic interaction between the two drugs in these genetic backgrounds. In contrast, for strains NCYC and Y9, no distinct 10-fold increase or decrease in growth was observed; consequently, their responses were classified as additive. This suggested that, in these genetic backgrounds, ibuprofen did not significantly modulate the efficacy of caspofungin.

When the drug combination was evaluated in CSM media (Fig. 2D), a substantial reduction in growth was observed across the strains in the combination plates. This effect was attributed to the presence of ibuprofen, the impact of which appeared to be amplified by the use of minimal CSM media, known to be metabolically demanding for cells, potentially exacerbating drug-induced stress. The caspofungin-resistant strains YJM789, 322134S, and DBVPG demonstrated a 100- to 10,000-fold reduction in growth when exposed to the combination, revealing potent synergistic interactions of the drugs. In contrast, no significant growth differences were observable for the caspofungin-sensitive strains NCYC, UWOP, and Y9, indicating that the drugs interacted additively in these genetic backgrounds.

The caspofungin-resistant strains YJM789, 322134S, and DBVPG consistently showed synergistic interactions between ibuprofen and caspofungin in both YPD and CSM media. In contrast, the caspofungin-sensitive strains NCYC and Y9 exhibited additive interaction across both media conditions. The caspofungin-sensitive strain UWOP exhibited contrasting behavior, showing a marginal synergistic interaction in YPD and an additive interaction in CSM media.

### 2.4. Assay type and concentration dependence of ibuprofen-clotrimazole interaction profile

To complement the semi-quantitative spot dilution assays, we performed quantitative broth dilution checkerboard assays to characterize ibuprofen–antifungal interaction patterns across the yeast strains. The most commonly selected reference models for evaluating drug interaction in a checkerboard assay are Loewe’s additivity model and the Bliss independence model (Ianevski et al., 2022). However, both models come with assumptions regarding the drug’s dose-response profiles or mechanisms. For instance, Loewe’s additivity is based on the assumption that the drugs exhibit similar dose-response curves (Lederer et al., 2018), which is often not the case with many drug combinations. In contrast, the Bliss independence model is grounded in the multiplicative survival principle, positing that the drugs should target independent pathways that do not share any mechanistic connections beyond the response outcome (Liu et al., 2018). It has been observed that for moderately effective drugs, the Bliss model anticipates a substantial increase in effect beyond that of the single agents to qualify as synergy (Wooten et al., 2021). Given that we were dealing with repurposed drugs that have not been well-studied for their targets in yeast and produced minimal effects independently, applying the Bliss reference model might skew the results towards antagonism. It is important to note that synergy can arise from drug combinations that address resistance mechanisms and does not necessarily require targeting other pathways within the cell (Zhu et al., 2023).

Other newly developed models, such as Zero Interaction Potency (ZIP), combine features of both Loewe’s additivity model and the Bliss independence model, yet each has its own limitations and assumptions (Yadav et al., 2015). Given these uncertainties, we opted to apply the simple reference model of the highest single agent (HSA) (Duarte & Vale, 2022) for drug interaction classification. We also explored synergy plots based on the other reference models, Bliss, Loewe, and ZIP, for both the drug combinations (Fig. S1, S2). It was evident that both the Bliss and ZIP models yielded similar results, with comparable interaction patterns across the drug spectrum. In contrast, Loewe’s additivity model produced a different interaction pattern, which could be attributed to its underlying assumptions.

In the liquid checkerboard assay, clinical strain YJM789 showed synergistic interactions between ibuprofen and clotrimazole, yielding a synergy score of +22.1 (Fig. 3A). This observation was consistent with results from the YPD solid agar assays. Conversely, strain 322134S showed divergent behavior across media types; while antagonistic effects were observed on solid agar, the liquid checkerboard assay revealed a synergistic interaction with a score of +16.1 (Fig. 3B). Although 322134S was a flocculating strain, experimental consistency was maintained by ensuring all wells were thoroughly mixed before optical density measurements.

**Figure 3:**
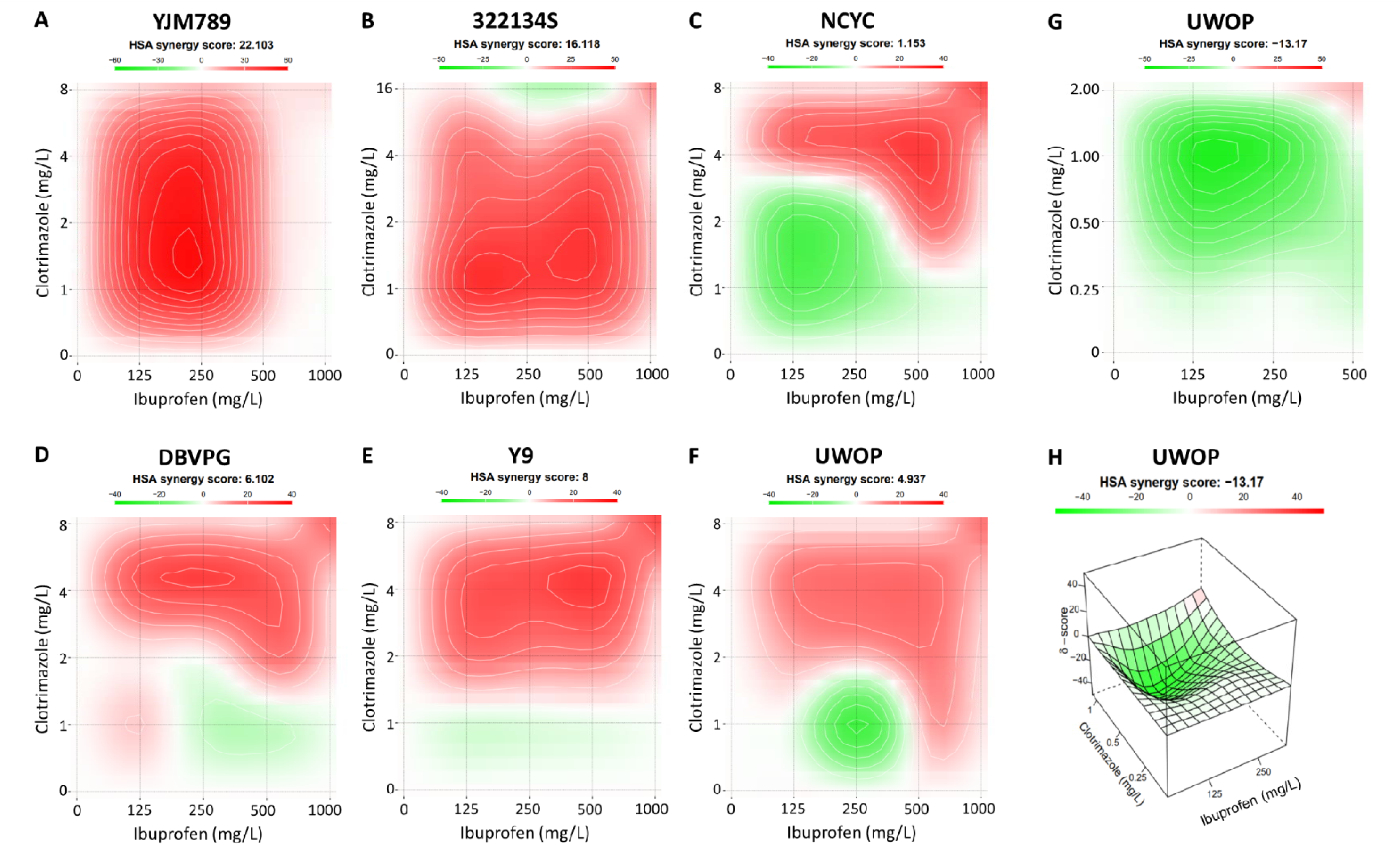
Dose-dependent drug interactions of ibuprofen and clotrimazole. (A-F) Synergy maps showing dose-dependent interactions between ibuprofen and clotrimazole across six S. cerevisiae strains. The x-axis represents the concentration of ibuprofen (mg/L), and the y-axis represents the concentration of clotrimazole (mg/L). 2D plot (G) and 3D plot (H) of the synergy scores for the UWOP strain tested with a narrow range of clotrimazole and ibuprofen. The color scale indicates the synergy score: red signifies synergy (HSA synergy score > 0), and green indicates antagonism (HSA synergy score < 0). The overall HSA synergy score for each strain is provided at the top of each plot.

For strain NCYC, the drug combination showed antagonism in the solid media assay but additivity in the liquid checkerboard assay, with a synergy score of +1.2. However, the synergy plots revealed distinct dose-dependent interaction patterns (Fig. 3C). Specifically, the combination demonstrated antagonism at lower concentrations of both agents (minimum synergy score: -24.9), whereas synergistic interactions emerged at higher dosage levels (maximum synergy score: +27.8). Similarly, strain DBVPG displayed antagonism in the YPD spot assay but an additive profile in the checkerboard assay, with a synergy score of +6.1 (Fig. 3D). Closer examination of the interaction landscape indicated that ibuprofen remained largely ineffective at lower concentrations of clotrimazole, regardless of the ibuprofen dosage. However, at elevated clotrimazole concentrations, ibuprofen demonstrated a clear synergistic effect, enhancing the overall inhibitory activity (maximum synergy score: +26.8).

Strain Y9 demonstrated antagonistic behavior of ibuprofen-clotrimazole in the YPD sold media assay, while the checkerboard assay demonstrated an additive global interaction (synergy score: +8.0); however, synergistic interactions were consistently observed within the combination space characterized by higher clotrimazole concentrations (Fig. 3E).

Strain UWOP revealed an antagonistic interaction of the drugs in the YPD agar assay and an additive interaction overall in the checkerboard assay (synergy score: +4.9). However, a more detailed analysis of the interaction landscape revealed significant spatial heterogeneity (Fig. 3F). While synergistic behavior was observed at higher clotrimazole concentrations (maximum synergy score: +18.4), a localized region of potent antagonism was identified, centered at 250 mg/L ibuprofen and 1 mg/L clotrimazole, where the local synergy score reached -26.3. This finding corroborated the antagonism observed in the YPD spot assays at 250 mg/L ibuprofen and 0.25 mg/L clotrimazole. To validate this concentration-specific antagonism, a targeted checkerboard assay was performed using a narrow range of clotrimazole concentrations (0-2 mg/L). This targeted concentration assay confirmed the antagonistic relationship, yielding a synergy score of -13.2 (Fig. 3G, H). These results showed that, for the ibuprofen-clotrimazole combination, the outcome of synergy or antagonism was not only dependent on the growth media and genetic background but also influenced by the specific assay conditions. Furthermore, in certain genetic backgrounds, such as the UWOP strain, the interaction was found to be modulated by specific drug dosages. These dose-dependent shifts introduced an additional layer of complexity to the phenotypic response, demonstrating that the strain’s response to a drug combination was a complex phenomenon.

### 2.5. Genotype-driven variability in ibuprofen-caspofungin interaction independent of assay format and concentration

YPD liquid checkerboard assay for the ibuprofen-caspofungin combination revealed that the caspofungin-resistant strains YJM789, 322134S, and DBVPG consistently exhibited synergistic interactions with synergy scores of +25.2, +18.5, and +13.5, respectively (Fig. 4A, B, and D). Notably, the synergistic effect in these three isolates was uniform across the entire tested concentration range, with no evidence of dose-specific or localized interaction shifts, and corroborated the observation from the YPD agar assay. The checkerboard assay of the caspofungin-sensitive strains NCYC, UWOP, and Y9 had synergy scores of +4.6, -0.2, and -2.0, respectively (Fig. 4C, E, F). These values indicated an overall additive interaction (i.e., drug combination had a synergy score value between -10 and +10). These findings are particularly noteworthy as there was a lack of concordance with previous results; specifically, strains UWOP and Y9 had previously demonstrated distinct synergistic and antagonistic profiles in solid agar assays, respectively. The transition from these varied interactions on solid media to a predominantly additive effect in the liquid checkerboard assay suggests that the physical state of the growth medium might have influenced the pharmacological dynamics of the ibuprofen-caspofungin combination.

**Figure 4:**
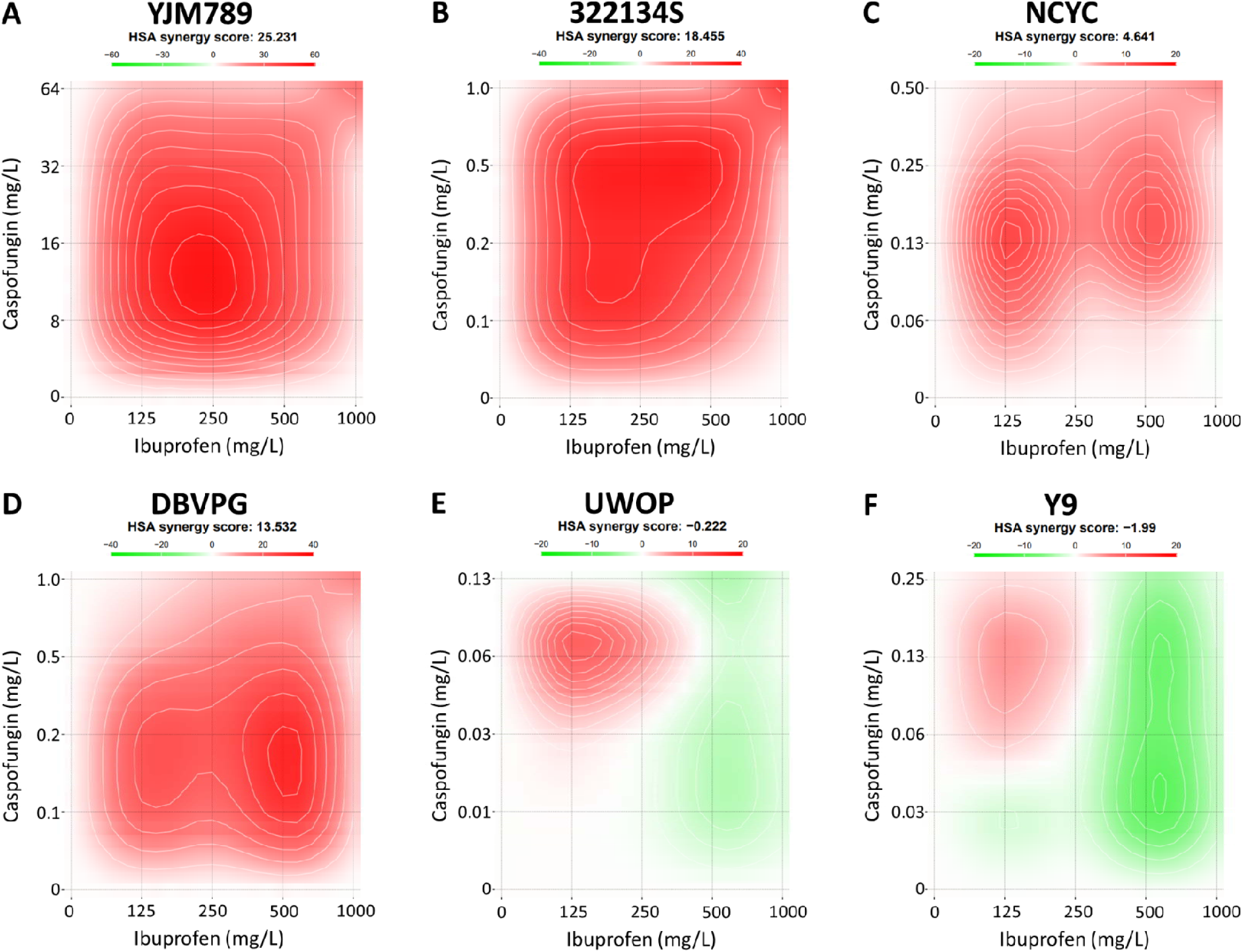
Dose-dependent drug interactions of ibuprofen and caspofungin. (A-F) Synergy maps showing dose-dependent interactions between ibuprofen and caspofungin across six S. cerevisiae strains. The x-axis represents the concentration of ibuprofen (mg/L), and the y-axis represents the concentration of caspofungin (mg/L). The color scale indicates the synergy score: red signifies synergy (HSA synergy score > 0), and green indicates antagonism (HSA synergy score < 0). The overall HSA synergy score for each strain is provided at the top of each plot.

## 3. Discussion

Characterizing drug interactions remains challenging due to ongoing debates regarding which reference models (such as HSA, Loewe’s additivity, Bliss independence, or ZIP) best capture true synergy. This is exemplified by the rapamycin-caspofungin combination, where the Loewe-based FICI method indicates additivity while the Bliss-based response surface methodology suggests synergy (Lefranc et al., 2024). The biological complexity of drug interactions, whether synergistic or antagonistic, goes beyond mathematical discrepancies. These models often fail to account for the underlying mechanistic pathways that determine outcomes.

While previous studies primarily report synergy between ibuprofen and azoles, often attributing this to ibuprofen’s ability to block efflux pumps in resistant clinical strains. Our results demonstrate a significant potential for antagonism between ibuprofen and azole. This variability is underscored by the clinical strains used in this study. While YJM789 exhibited synergy, strain 322134S showed clear antagonism, suggesting that ibuprofen may interfere with azole efficacy in strains where resistance is not efflux-mediated. Furthermore, the transition from synergy to additivity observed in YJM789 when moved from rich to minimal media may be driven by metabolic constraints. In CSM media, high energy demand for nutrient biosynthesis is likely to restrict the cellular energy budget (Fujiwara et al., 2025) and limit the ATP available for efflux pump activity, thereby reducing the synergistic impact of ibuprofen on efflux pump inhibition. Furthermore, it is noteworthy that the clinical strains exhibited non-linear dose responses to clotrimazole. For instance, YJM789 displayed similar growth-inhibited phenotypes at both 0.25 mg/L and 1 mg/L of clotrimazole. Such biphasic or non-monotonic dose-response curves can significantly complicate the interpretation of drug combinations, necessitating detailed mechanistic studies to understand how these dynamics modulate overall interaction patterns.

In YPD solid media assays, antagonistic interactions between ibuprofen and azoles were consistently observed in wild and fermentation isolates, whereas liquid checkerboard assays showed concentration-specific effects. This contrasting result suggests that drug interaction outcomes are significantly modulated by the experimental platform, likely driven by differences in oxygen availability, drug exposure, and metabolic heterogeneity between solid and liquid cultures. Standard synergy classification methods, such as the Loewe-based FICI, fail to capture the concentration-dependent interactions we observed in half of the tested strains. While previous studies on *C. albicans* reported predominant synergy, our results in *S. cerevisiae* revealed antagonism in five of six strains in rich media, highlighting the influence of genetic diversity and potential species-specific metabolic differences. By using a diverse panel of wild isolates rather than a limited set of clinical strains, we captured interaction patterns that are typically overlooked in traditional antifungal screening. These naive strains, not exposed to clinical drug stress, provide a unique baseline for evaluating the full spectrum of potential responses to drug combinations. Future research must validate this context-dependent framework in clinical isolates to determine if these antagonistic effects translate to human pathogens.

In contrast, the less-explored ibuprofen-caspofungin combination appears more predictable; a strain’s response to the drug pair generally correlates with its baseline resistance to caspofungin alone (Table 3). While caspofungin directly inhibits 1,3-β-D-glucan synthase at the plasma membrane, ibuprofen’s reported ability to alter lipid membrane composition and induce reactive oxygen species (ROS) (Babaei et al., 2024) may act orthogonally to the glucan synthesis pathway. This mechanistic divergence likely explains why caspofungin-resistant strains exhibit potent synergy, suggesting that ibuprofen potentiates antifungal efficacy through alternative, novel pathways. To validate these findings, future research should quantify the roles of ROS and lipid membrane fluidity in ibuprofen-resistant strains.

**Table 3:** Overall results of the ibuprofen-antifungal studies.

| Strains | Ibuprofen-Clotrimazole |  |  | Ibuprofen-Caspofungin |  |  |
| --- | --- | --- | --- | --- | --- | --- |
|  | YPD_Liq | YPD_solid | CSM_solid | YPD_Liq | YPD_solid | CSM_solid |
| YJM789 | Synergy | Synergy | Additive | Synergy | Synergy | Synergy |
| 322134S | Synergy | Antagonism | Additive | Synergy | Synergy | Synergy |
| NCYC | Additive | Antagonism | Additive | Additive | Additive | Additive |
| DBVPG | Additive | Antagonism | Additive | Synergy | Synergy | Synergy |
| UWOP | Additive | Antagonism | Additive | Additive | Synergy | Additive |
| Y9 | Additive | Antagonism | Antagonism | Additive | Additive | Additive |

Our results demonstrate that the response to repurposed drug combinations is not a static phenomenon, but a complex phenotype shaped by genetic background and environmental context. Further, pairing repurposed drugs with different classes of conventional antifungals reveals a broader spectrum of interaction mechanisms that might otherwise remain undiscovered. These findings highlight the need for careful, comprehensive strain-level optimization before the clinical deployment of a drug combination to improve efficacy.

To effectively explore repurposed drug combinations to capture the full biological complexity of these pairings, it is crucial to conduct in vitro experiments that mimic the nutrient profiles and microenvironments of specific infection sites and to test within established human therapeutic windows to generate results that more closely align with patient responses. While this study demonstrates the potential of *S. cerevisiae* as a proof-of-concept, the use of non-pathogenic species is a critical limitation, necessitating further research on clinical isolates to validate and translate these findings to pathogenic species. Prioritizing the characterization of molecular pathways governing genotype-environment interactions is essential, as it will facilitate the translation of innovative drug combinations into impactful clinical therapies, ultimately improving patient care.

## 4. Materials and Methods

### 4.1. Strains, drugs, and growth conditions

Six *S. cerevisiae* haploid yeast strains, YJM789 (clinical isolate), 322134S (clinical isolate), NCYC110 (fermentation isolate; hereafter NCYC), DBVPG1373 (wild isolate; hereafter DBVPG), UWOPS03-461.4 (wild isolate; hereafter UWOP), and Y9 (fermentation isolate), were selected from the Saccharomyces Genome Resequencing Project (SGRP) collection (Liti et al., 2009) to represent a diversity of geographic origins and ecological niches (Table 1). Strains were grown on yeast peptone dextrose medium (YPD; 1% Yeast extract, 2% Peptone, and 2% Glucose) or complete synthetic medium (CSM; 0.14% Yeast Nitrogen Base, 0.50% Ammonium sulfate, 0.077% Complete Supplement Mixture, and 2% Glucose) adjusted to pH 5.8 with 2.5 M NaOH. The media were supplemented with 20 g/L agar for solid cultures, and broth medium was used for liquid cultures. The antifungal agents used in this study were clotrimazole (Sigma-Aldrich, Cat# C6019), caspofungin (Cayman Chemical, Cat# 15923), and the repurposed non-antifungal drug, ibuprofen (TCI Chemicals, Cat# I0415). Stock solutions of each drug were prepared in sterile dimethyl sulfoxide (DMSO, HiMedia, Cat# MB508)

### 4.2. Spot dilution assay

An overnight saturated culture of each yeast strain was prepared by inoculating a single colony into 5 mL of YPD broth and incubating at 30°C. The optical density (OD) of each culture was measured (Amersham Biosciences Ultrospec 2100 Pro), and the cell suspensions were normalized to an OD600 of 4.0, corresponding to approximately 10^8^ cells/mL. The normalized cell suspension was then subjected to five consecutive 10-fold serial dilutions. Five microliters from each dilution were spotted onto YPD or CSM agar plates. Control plates lacked drugs, while test plates contained one or both drugs at various concentrations. All plates were incubated at 30°C for three days and photographed at 24, 48, and 72 hours. Observations at 48 hours were used to determine drug interaction patterns, while observations at 24 and 72 hours were used to balance the lag phase and potential adaptation effects. All experiments were performed in triplicate. Growth was defined by the number of visible spots.

### 4.3. Checkerboard assay

The drug interaction between antifungal and non-antifungal agents was assessed using the checkerboard assay. The drug concentration range for each strain was first optimized to ensure optimal results from the checkerboard assay. The minimum inhibitory concentration (MIC), defined as greater than 90% growth inhibition (Lefranc et al., 2024), was determined for each drug-strain combination, as listed in Table 2. We performed 5×5 checkerboard assays in a 96-well microtiter plate according to a defined protocol (Bellio et al., 2021). Briefly, isolates were inoculated at 10^6^ cells/mL into YPD medium. The broth contained varying concentrations of two drugs, with one drug concentration exponentially varying along the abscissa and the other along the ordinate(Bellio et al., 2021). For all tested combinations, the concentration range spanned from 0 to the MIC to comprehensively explore the drug interaction space. To investigate the concentration-specific interaction patterns observed in the UWOP strain for ibuprofen-clotrimazole combination, a targeted checkerboard assay was performed using a narrowed clotrimazole range (0-2 mg/L) in combination with ibuprofen (0-500 mg/L). Following inoculation, plates were incubated in a shaking incubator at 30°C. Cell growth was then quantified using a microplate reader (Agilent BioTek Epoch2NS) at an OD600 after 48 hours of incubation. All the experiments were done in quadruplicate.

### 4.4. Drug interaction patterns

The percentage of viable cells was calculated and utilized as input for SynergyFinder (Ianevski et al., 2022). Based on the selected reference model, synergy scores were generated to classify drug interactions as synergistic, additive, or antagonistic. While SynergyFinder supports multiple reference models, such as Loewe’s additivity and Bliss independence, and Zero Interaction Potency (ZIP), all these complex models carry specific assumptions regarding drug dose-response profiles and mechanisms of action. Previous research has indicated that strain-specific dose-response curves can be highly complex and unique, often eluding the predictive capacity of these models (Schmidlin et al., 2025). Given the uncertainty surrounding the actual mechanism of ibuprofen in yeast and the varying dose-response curves when compared to clotrimazole and caspofungin, we opted to utilize the simple reference model of the highest single agent (HSA) (Duarte & Vale, 2022) for drug interaction classification. The web-based tool was executed using the default parameters specified in the user manual. Consistent with the software’s guidelines, an antagonistic interaction was defined as a synergy score below -10, a synergistic interaction as a score above +10, and an additive interaction as a score between -10 and +10. Additionally, SynergyFinder generated synergy maps to visualize the interaction landscape, highlighting both synergistic and antagonistic dose combinations. In these 2D synergy maps, the axes represented drug concentrations, while the 3D maps incorporated the synergy score as the third dimension.

## Acknowledgements

We acknowledge discussions and support from all members of the Systems Genetics Lab, the Centre for Integrative Biology and Systems Medicine (IBSE), and the Wadhwani School of Data Science and Artificial Intelligence (WSAI). A.P. was supported by Prime Ministers’ Research Fellowship, Department of Education, Govt. of India (SB/2223/1564/BT/PMRF/008752). A.P. also acknowledges a travel fellowship from the IBSE grant to H.S. (BIO/1819/304/ALUM/KARH). H.S. acknowledges funding support from CEFIPRA (IFC/6103-2/2019/500).

Conceptualization: H.S.; Investigation, Methodology, Analysis, Visualization: A.P. Supervision: H.S.; Writing-original draft: A.P., Writing-review and editing: A.P., H.S.

## Data availability statement

All data generated during this study are included in the article and the supplementary information.

## Supplementary Information

**Figure S1:**
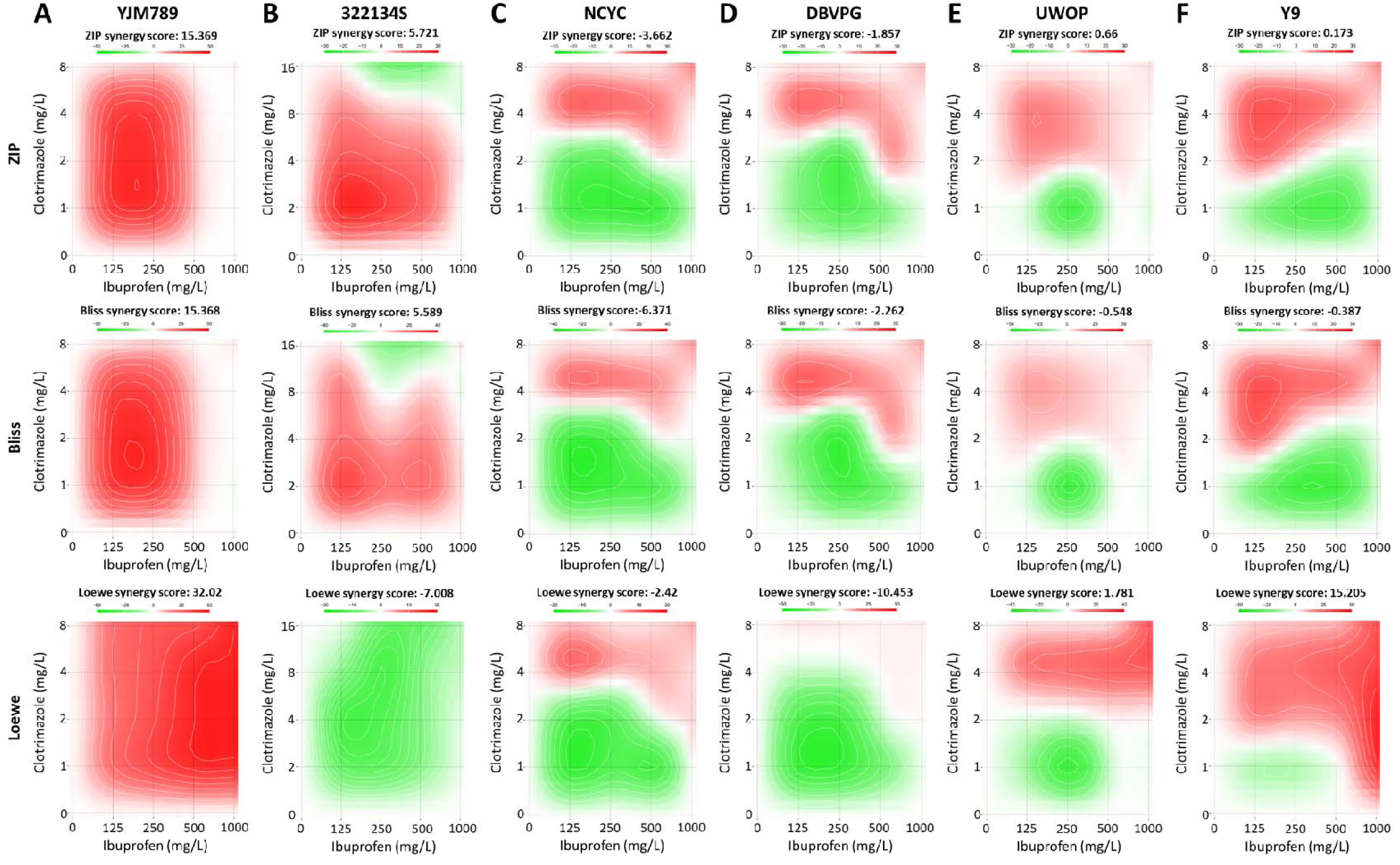
Synergy landscapes for Ibuprofen-Clotrimazole combinations across six S. cerevisiae strains. The synergy maps illustrate drug interaction scores, where each column represents a single yeast strain (A-F) and each row corresponds to a specific reference model: Zero Interaction Potency (ZIP), Bliss Independence (Bliss), and Loewe Additivity (Loewe). Red regions indicate synergistic inhibition, while green regions signify antagonistic interactions across the concentration gradients of both drugs.

**Figure S2:**
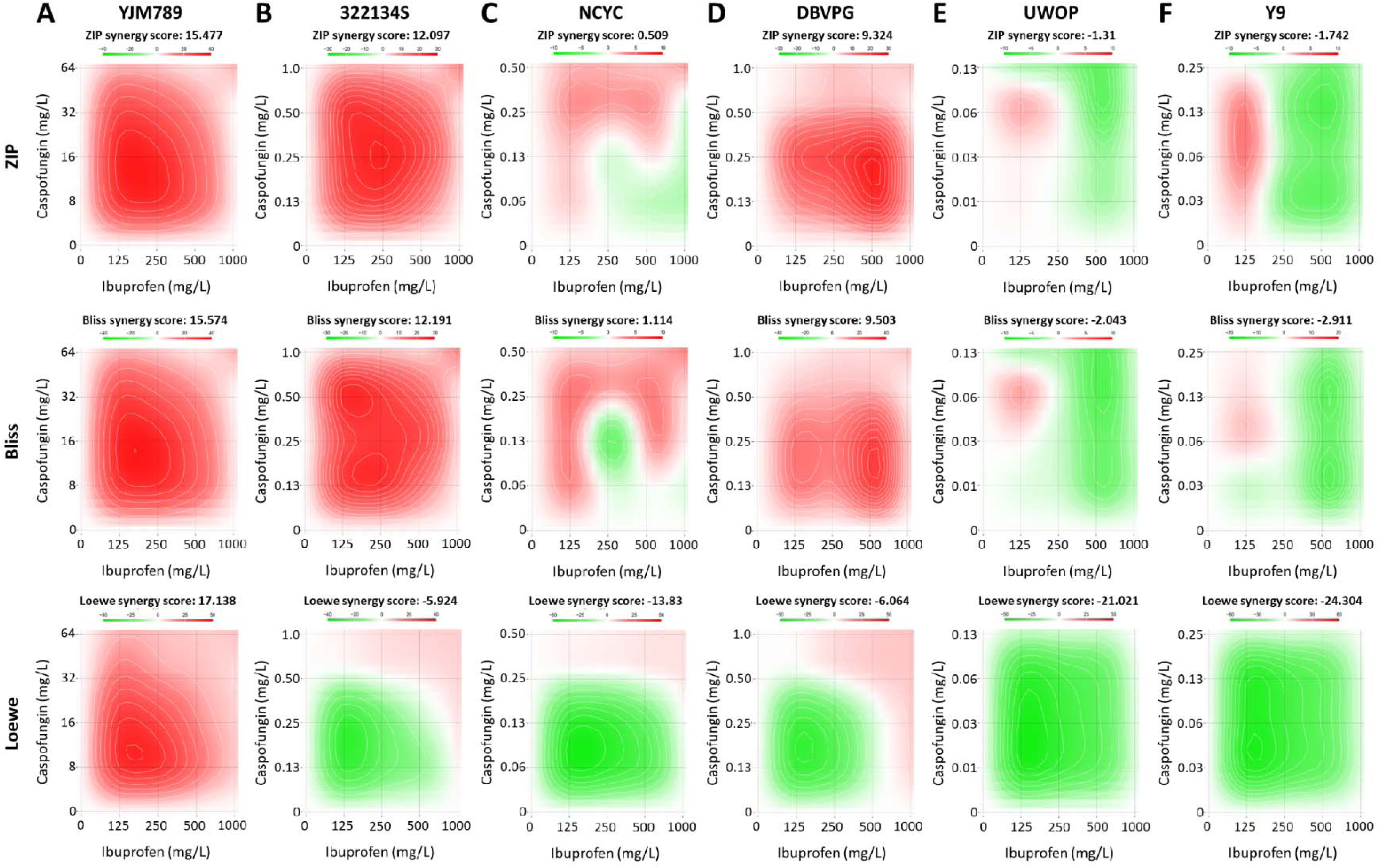
Synergy landscapes for Ibuprofen-Caspofungin combinations across six S. cerevisiae strains. The synergy maps illustrate drug interaction scores, where each column represents a single yeast strain (A-F) and each row corresponds to a specific reference model: Zero Interaction Potency (ZIP), Bliss Independence (Bliss), and Loewe Additivity (Loewe). Red regions indicate synergistic inhibition, while green regions signify antagonistic interactions across the concentration gradients of both drugs.

## Notes

### Competing Interest Statement

The authors have declared no competing interest.

